# Periods of environmental sensitivity couple larval behavior and development

**DOI:** 10.1101/2023.08.04.552015

**Authors:** Denis F. Faerberg, Erin Z. Aprison, Ilya Ruvinsky

## Abstract

The typical life cycle in most animal phyla includes a larval period that bridges embryogenesis and adulthood^1^. Despite the great diversity of larval forms, all larvae grow, acquire adult morphology and function, while navigating their habitats to obtain resources necessary for development. How larval development is coordinated with behavior remains substantially unclear. Here, we describe features of the iterative organization of larval stages that serve to assess the environment and procure resources prior to costly developmental commitments. We found that male-excreted pheromones accelerate^2-4^ the onset of adulthood in *C. elegans* hermaphrodites by coordinately advancing multiple developmental events and growth during the last larval stage. The larvae are sensitive to the accelerating male pheromones only at the end of the penultimate larval stage, just before the acceleration begins. Other larval stages also contain windows of sensitivity to environmental inputs. Importantly, behaviors associated with search and consumption of food are distinct between early and late portions of larval stages. We infer that each larval stage in *C. elegans* is subdivided into two epochs: A) global assessment of the environment to identify the most suitable patch and B) consumption of sufficient food and acquisition of salient information for developmental events in the next stage. We predict that in larvae of other species behavior is also divided into distinct epochs optimized either for assessing the habitat or obtaining the resources. Thus, a major role of larval behavior is to coordinate the orderly progression of development in variable environments.

“Life is what happens to you when you’re busy making other plans.”

Allen Saunders (popularized by John Lennon)

## Results and Discussion

### Male-excreted signals accelerate reproductive development and larval growth in *C. elegans* hermaphrodites

*C. elegans* hermaphrodite larvae accelerate development in the presence of male-excreted signals^2-4^. Evidence to support this claim largely comes from the earlier acquisition of adult vulva morphology (staging shown in Figure S1A) in hermaphrodites exposed to male signals (Figure 1A, B). We tested whether development of other aspects of the reproductive system were similarly accelerated. On male-conditioned plates (MCPs), the appearance of morphologically defined oocytes (Figure 1C, S2A) and embryos in the uterus (Figure 1D, S2B) were advanced by approximately the same two hours as the appearance of adult vulva morphology. The substantial difference in the time-to-adulthood between the slowest- and the fastest-developing worms (>6 hours; Figure 1A, B) reflects the heterogeneity of rates of larval developmental, as observed previously^5-8^.

**Figure 1.**
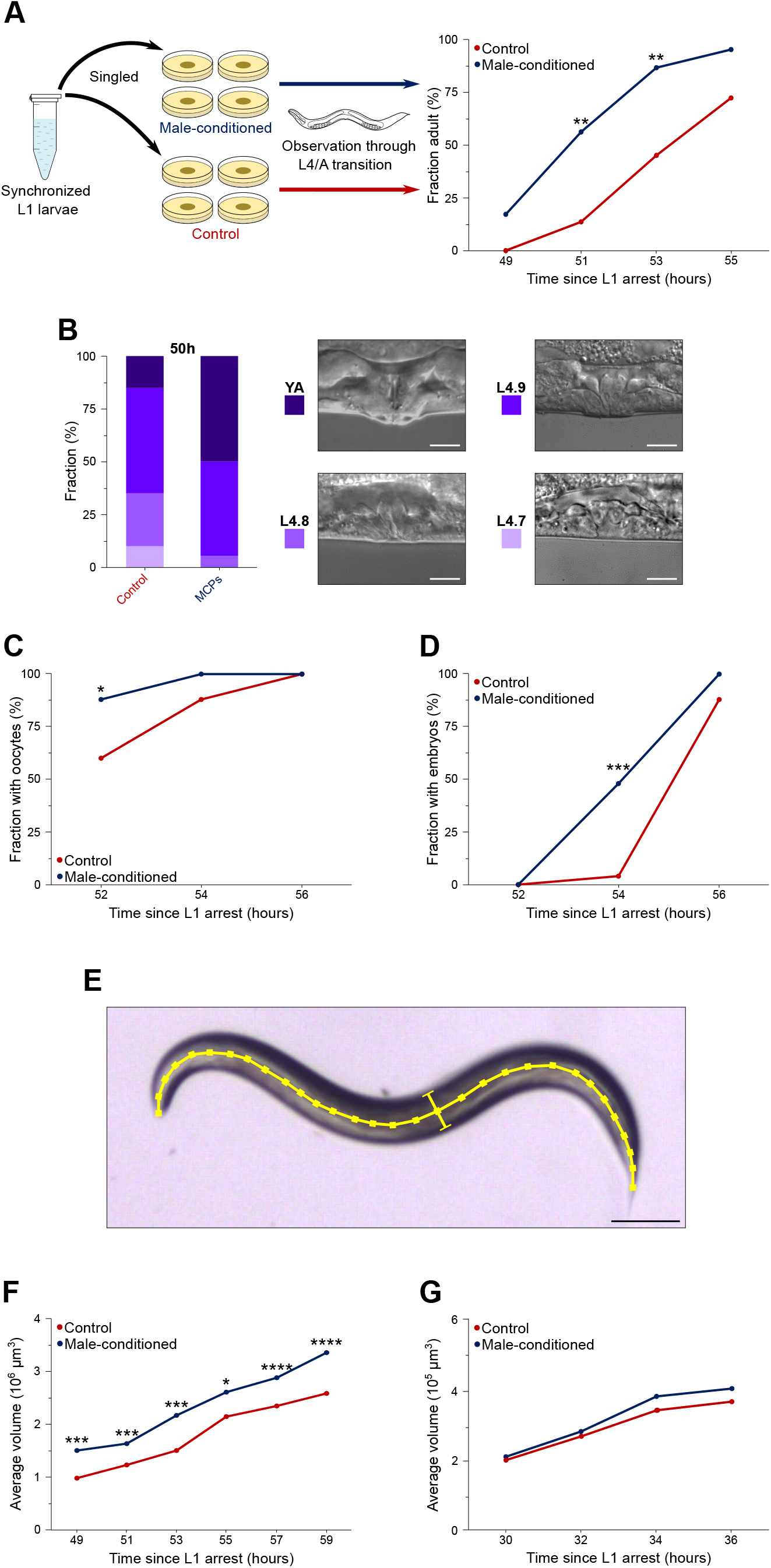
Male-excreted signals accelerate late larval development in *C. elegans* hermaphrodites. Synchronized L1 larvae were individually grown on plates conditioned by live male habitation (male-conditioned, or MCPs; blue) or on control plates (control; red). Individuals on MCPs achieved adult vulva morphology ∼1.5-2 hours earlier. **(B)** Distributions of morphological vulva substages (as defined on the right; also Figure S1A) on control and male-conditioned plates. Scale bars = 10μm. **(C)** Frequency of individuals with at least one oocyte spanning the gonad lumen. **(D)** Frequency of individuals with at least one embryo in the uterus. **(E)** Sample worm image used to estimate body volume. Yellow lines indicate the midline (length) and cross-section (width) for ImageJ processing. Scale bar = 100μm. **(F)** Average volume of hermaphrodites in early adulthood estimated from length and width measurements. **(G)** Average volume of hermaphrodites in the second half of the L3 larval stage. See Table S1 for sample sizes and statistical analyses.

During larval development *C. elegans* hermaphrodites increase in volume almost 100-fold^6,9,10^. It is possible that in the presence of male signals hermaphrodites experience earlier maturation of the reproductive systems while the growth schedule remains unaltered. We found that by the time of larval-sto-adult transition, worms on MCPs were consistently larger than the controls (Figure 1E, F). The accelerated growth was isomorphic (Figure S2C) and advanced by approximately the same two hours as the accelerated development of the reproductive system. The divergence in size between the control and MCP worms became evident around 34 hours into larval development (throughout this study time refers to hours from release from the L1 arrest); this time corresponds to the late L3 stage (Figure 1G, S2D, and text below).

### Systemic acceleration of development around the L3-to-L4 transition

The classical detailed description of the entire post-embryonic cell lineage in *C. elegans*^11^ allows a more precise determination of which developmental processes in hermaphrodite larvae accelerate on MCP, and when. The H and V blast cells and their progeny undergo cell divisions at the beginning of L2, L3, and L4 stages (Figure 2A, S2B) to give rise to the hypodermal seam cells; adhesion occurs later in the stages^12^. Whereas no difference was evident around 26-28 hours, larvae on MCPs have started to accelerate as early as ∼36-38 hours. Because cell division events are closely tied to molting episodes, the fractions of worms exhibiting a particular number of cells can serve as a measure of developmental progression. At 26 hours, ∼30% of worms had an L3-specific number of cells, indicating that in our hands the fastest larvae started this stage around 24 hours. Using similar logic, the fastest-developing larvae reached the L4 stage just prior to 36 hours when ∼13% of worms had an L4-chracteristic number of seam cells.

**Figure 2.**
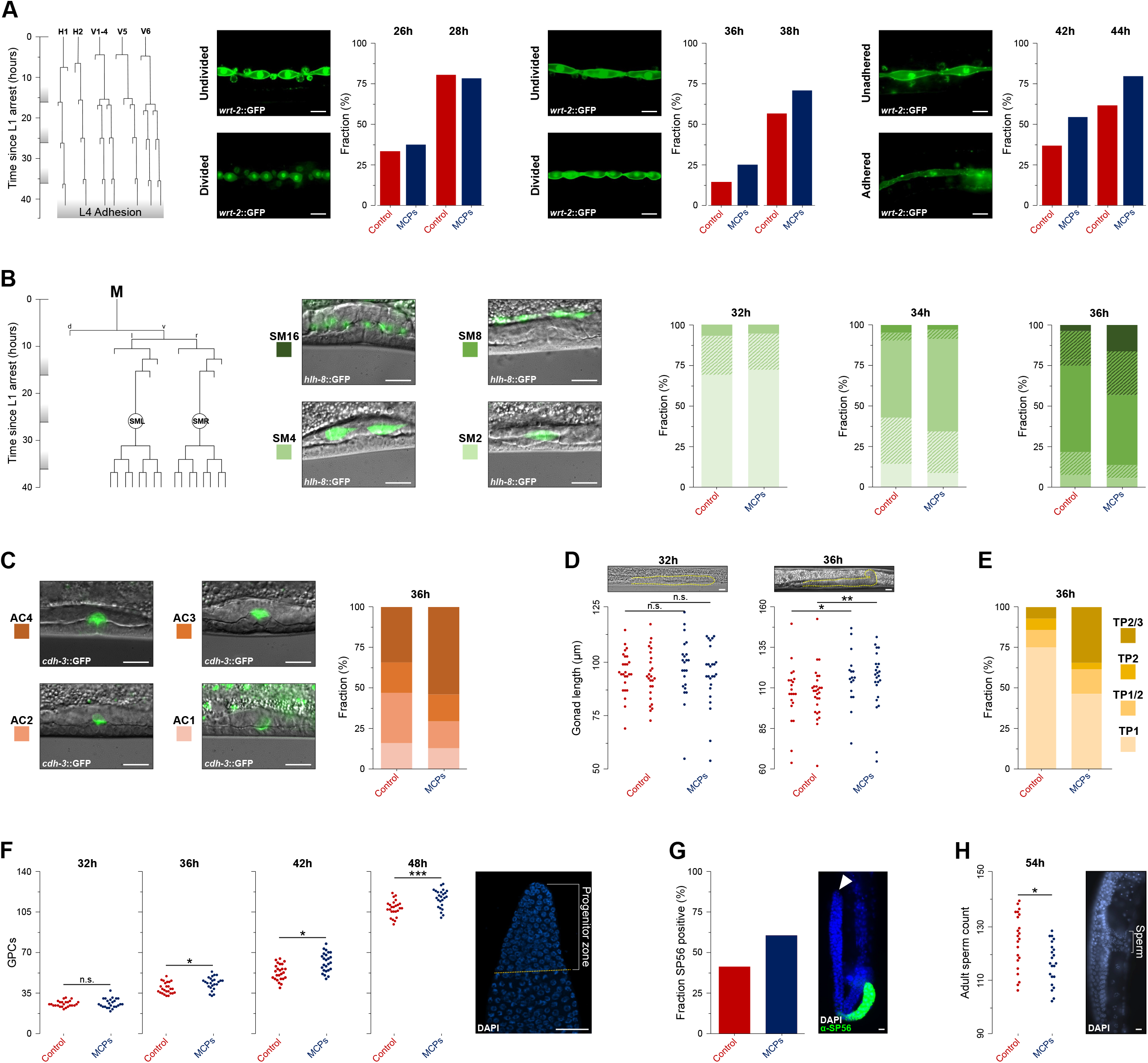
Male signals systemically accelerate larval development in hermaphrodites around the L3/L4 transition. (A) Seam cells divide after molts (grey boxes along the time axis) and undergo subsequent adhesion events at times indicated in the lineage diagram (derived from^11^). Bar plots show fractions of worms with initiated L3 seam cell division (left panel, 26-28h), L4 division (middle panel, 36-38h), and L4 adhesion (right panel, 42-44h). Additional information is in Figure S1B. **(B)** Sex myoblast cells (M.vlpaa and M.vrpaa) undergo three rounds of division around the L3/L4 transition (derived from^11^). Bar plots show fractions of larvae with indicated numbers of cells. Solid portions indicate individuals with completed divisions. Dashed portions indicate individuals with an initiated, but unfinished division round. Additional information is in Figure S1C. **(C)** The anchor cell attaches to P6 daughter cells and initiates invagination around the L3/L4 transition. The bar plot shows frequencies of each anchor cell “stage”. Additional information is in Figure S1D. **(D)** Gonad lengths measured at midline (gonad outlines shown in yellow in images above). In each group, the left data series represents anterior arms, the right is posterior arms. **(E)** Quantification of gonad turning which occurs around the L3/L4 transition. The bar plot shows frequencies of each phase of gonadal turning. Additional information is Figure S1E. **(F)** Numbers of germline precursor cells. Dashed yellow line in the inset indicates the boundary of the Progenitor Zone. **(G)** Fraction of gonad arms expressing SP56, an early marker of sperm differentiation. White arrowhead marks the distal end of the gonad. **(H)** Number of self-sperm in adult hermaphrodites. In panels **D, F**, and **H**, each dot represents measurements/counts from one gonad arm. All scale bars = 10μm. See Table S1 for sample sizes and statistical analyses.

We next examined the timing of the sex myoblast divisions (Figure S1C). These cells are the progeny of the M blast cell, which comes from a different founder cell (MS) than the H and V blast cells (AB)^13^. We saw no difference in division timing on control plates vs. MCPs at 32 and 34 hours, the first evidence of divergence being detectable at 36 hours (Figure 2B). Given the timing of divisions in the sex myoblast lineage^11^, this marker places the L3-to-L4 transition around 34-36 hours, same as the estimate from the hypodermal seam cells (Figure 2A). Another descendent of the MS lineage, the vulva anchor cell, undergoes a series of morphological transformations (Figure S1D) between L3 and early L4^14^. The first sign of acceleration of the sequence of morphological transformations of the anchor cell on MCPs was evident around 36 hours (Figure 2C).

During late larval development, the hermaphrodite gonad changes considerably (Figure S1E)^15^. The first evidence of accelerated gonadal growth was detected around the L3-to-L4 transition at 36 hours (Figure 2D). In addition to growth, the gonad undergoes characteristic morphological changes^15-17^.

Accelerated turning became apparent by approximately 36 hours (Figure 2E). Starting in mid to late L3, the population of germline progenitor cells rapidly expands^18^. On MCPs this process was accelerated starting around 36 hours (Figure 2F). We note that this accelerated expansion of the larval germline is qualitatively different than the increased germline proliferation in the presence of the male pheromone ascr#10, because the latter only affects egg-laying adults^19,20^. The antibody that recognizes minor sperm proteins (SP56) is an early marker of sperm differentiation^21^, a process that begins early in L4^22^. Using this marker, we found that sperm differentiation was accelerated on MCPs (Figure 2G). The switch from spermatogenesis to oogenesis, that normally takes place in L4^23^, also appears to occur earlier because we observed fewer sperm cells in adult hermaphrodites (Figure 2H). Incidentally, earlier termination of spermatogenesis is consistent with the overall physiological response of the hermaphrodite reproductive system to the signals that indicate the presence of males – to redirect resources into oogenesis^20,24^.

Taken together, the data presented in Figures 1 and 2 argue that male-excreted signals systemically speed up growth as well as somatic and germline development in hermaphrodite larvae, as would be expected given the coupling between the growth rate and developmental tempo^6,25^. The acceleration starts at 34-36 hours, around the L3-to-L4 transition, and is thus mostly restricted to the last larval stage, consistent with previous findings^2-4^.

### Male signals influence commitment to the lethargus between L3 and L4 stages

In *C. elegans*, larval stages are separated by lethargus periods that typically last one to two hours^26^ and are characterized by sleep-like behavioral quiescence^27^. Therefore, we considered three possible, although not mutually exclusive hypotheses to account for developmental acceleration: A) earlier termination of the L3 stage, B) shorter lethargus, C) shorter L4. To distinguish between these possibilities, we monitored pharyngeal pumping in individual animals during the period of the L3-to-L4 transition.

Cessation of pumping is an indicator that worms are in lethargus^27,28^. A comparison of activity profiles on MCPs vs. control plates (Figure 3A) yielded five findings. First, contradicting Hypothesis A above, hermaphrodite larvae on MCPs exited L3 later than their counterparts on control plates (fraction of population in Figure 3B; age at first episode of quiescence in Figure 3C). Second, duration of quiescence immediately preceding the L4 stage was somewhat shorter on MCPs, a difference largely due to the reduction of long (>100 minutes) lethargus episodes (Figure 3D). Third, these two differences between activity schedules on MCPs and control plates cancelled each other since the onset of L4 was indistinguishable between these conditions (Figures 3B, S3A). Fourth, in this experiment, ∼1/2 of the larvae exited L3 around 34 hours and entered L4 around 36 hours (Figure 3B), nearly perfectly consistent with the estimates based on developmental markers (section above). Finally, a notable difference in activity around the L3-to-L4 transition was that many fewer MCP larvae showed “uncertain” entry into lethargus, that is, instances when an episode of inactivity was followed by the resumption of pumping and then another episode of quiescence before emerging as an L4 (Figure 3E).

**Figure 3.**
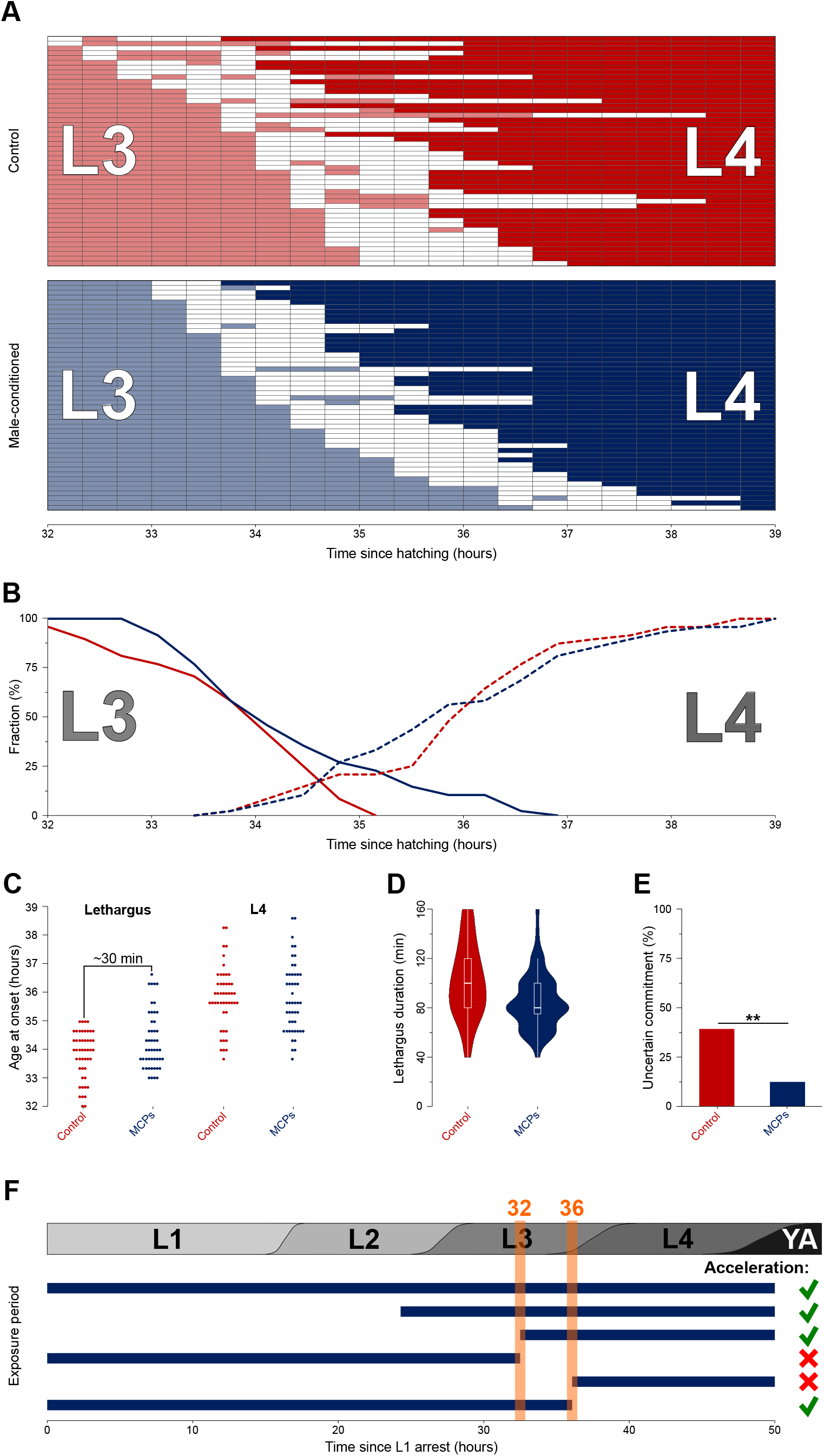
Male signals and developmental time. (A) Pharyngeal pumping activity, assayed at 20-minute intervals, of 48 control (red) and 48 MCP (blue) hermaphrodite larvae around the L3/L4 transition. The worms (each represented by one row) were sorted by the timing of the first observation period when pumping was not detected. See Figure S3A for an alternative plotting. Pale color = L3; white = non-pumping, presumed lethargus; intense color = L4. **(B)** Fractions of 48 individuals in L3 (solid lines) and L4 (dashed lines). **(C)** Age of worms at first entry into presumed L3/L4 lethargus (left) and at the onset of L4 (right). Each dot = one individual. Mean differences (MCPs – control) were 29.6 minutes for the entry into lethargus and -1.3 minutes for the onset of L4. **(D)** Larvae on MCPs spent less time in the L3/L4 lethargus (the averages differed by 15.8 minutes). **(E)** Fractions of 48 individuals with uncertain lethargus commitment. Entrance was judged as uncertain if a period of no pumping was followed by a period with pumping, but before the onset of L4. **(F)** Sensitivity window to male-excreted signals that induce developmental acceleration. Experiments (following the schematic in Figure 1A) with different exposure regimes. The grey bar above shows the approximate durations of larval stages. Sigmoidal transition boundaries represent inter-individual variability in developmental rates. Horizontal blue bars indicate the period of exposure to male-excreted signals. Green and red marks are acceleration and lack of acceleration, respectively. See Figure S3B for representative developmental plots. See Table S1 for sample sizes and statistical analyses.

Male-excreted signals prolonged the L3 stage, presumably to extend the feeding period, and increased the probability of irreversible commitment to lethargus on the first attempt. This view is corroborated by the comparison of growth curves. At 32, 34, and 36 hours, the MCP larvae were respectively 5%, 11%, and 10% larger than the paired controls (Figure 1G). The feeding fraction (number of episodes of feeding divided by total number of recorded episodes for all animals) was 6% greater on MCPs up to 34 hours and 13% greater between 34 and 36 hours (Figure 3A). Extended duration of feeding appears to account for the increased size of MCP worms.

### A narrow window of sensitivity to male-excreted signals during the late L3 stage

Because larvae on MCPs and the control plates entered L4 almost simultaneously (Figure 3B, C), the ∼2-hour acceleration (Figure 1A) likely occurred early in L4. We saw no developmental acceleration until∼36 hours (Figure 2), when ∼1/2 of the population was in L4. Whereas early in L4 the extent of acceleration was modest (in Figure 2A, worms on MCPs at 36 hours were substantially less advanced than the control at 38 hours), by mid-late L4 the MCP advantage increased to ∼2 hours (worms on MCPs at 42 hours almost equivalent to control at 44 hours, Figure 2A). The earlier onset of spermatogenesis (Figure 3G) and fewer mature sperm (Figure 3H) are also consistent with this view, because these processes take place early in L4^23^.

We transferred larval hermaphrodites between MCPs and control plates at different ages and for different periods of time to determine when they needed to experience male-excreted signals to accelerate growth and development. We found that a narrow time window between 32 and 36 hours, that is, from approximately mid/late L3 to the earliest L4 was required for the acceleration (Figure 3F, S3B). The strict tests of sufficiency of MCP exposure during this time for acceleration were inconclusive. This may be because exposure during other periods could contribute to acceleration and because it is technically difficult to narrow down the window of sensitivity to beyond the extent of temporal variability of development. By the L3-to-L4 transition, the fastest developing animals could be as much as 4-5 hours ahead of the slowest (Figure 3B, C).

### Coordination of larval development, behavior, and environmental sensitivity

Our results show that hermaphrodite larvae can follow one of two alternative scenarios during the L4 stage – the slower “standard” growth and development or the faster one that occurs in the presence of male-excreted signals (Figures 1, 2). For the acceleration to take place, exposure to male-excreted signals must occur during a narrow window in the late L3 stage, just prior to the onset of L4 (Figure 3F). The same critical time window was previously implicated in another paradigm. *C. elegans* hermaphrodites raised in isolation are smaller than their colony-raised counterparts; this growth defect could be rescued by transferring singled worms into groups of other hermaphrodites, but only until the onset of L4^29^.

Whether this growth phenotype is related to the one we identified in this study (Figure 1F) is currently unclear. The late L3 stage also appears to be the sensitive period when the male pheromones modulate synaptic transmission and, consequently, locomotion in hermaphrodites^30^.

There are parallels between the sensitive period in the late L3 and another point of potential bifurcation of developmental trajectories during *C. elegans* larval development. Under adverse conditions, larvae can opt out of reproductive development to become an alternative morph called dauer^31^. As is the case during the L3 decision, small-molecule pheromones are salient signals for dauer development^32,33^.

Another similarity is that the decision to enter dauer can only be made during the mid/late L1 stage, but not after the onset of L2^34^.

Generalizing the similarities between L1 and L3, we hypothesize that other developmental stages in *C. elegans* contain discrete periods of sensitivity to environmental signals (Figure 4A). There is evidence consistent with this view, although periods of sensitivity are less well defined than for L1 and L3. Late embryos sense their environment to select developmental trajectories in L1^35^. Experiences during the L2 stage impact developmental phenotypes in L3^36^. During a short window early in adulthood, just before the onset of egg laying, hermaphrodites are particularly sensitive to food and male pheromones that determine investment in and quality control of the oocytes^20^. Perception of social environment regulates the relative timing of somatic and germline development^37^.

**Figure 4.**
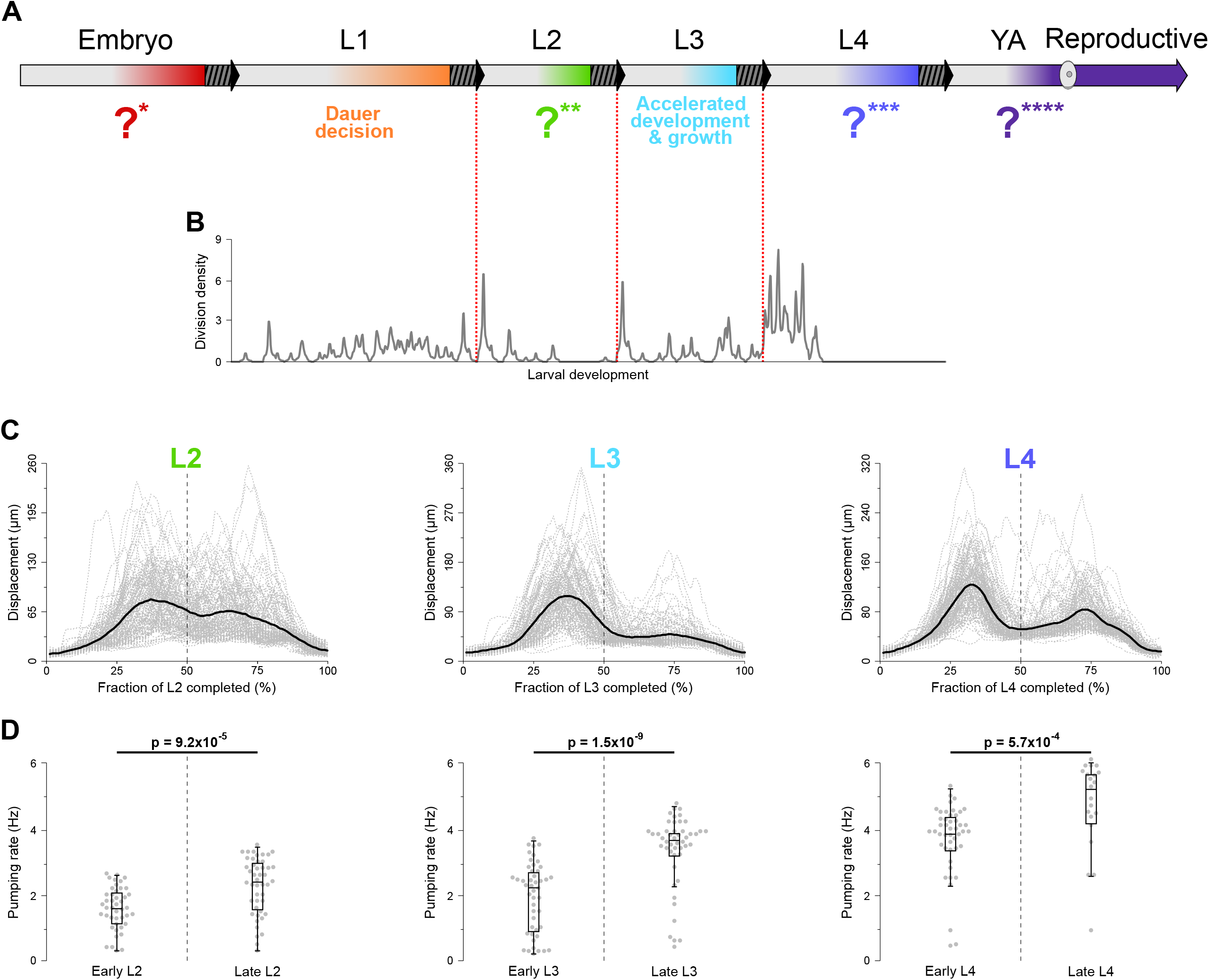
Coordination of larval development, behavior, and environmental sensitivity. (A) Schematic of *C. elegans* life history. Striped-grey blocks and arrows indicate transitions between stages (lethargus and ecdysis between larval stages). Color gradients represent periods of environmental sensitivity and commitment. **(B)** The density plot of cell divisions relative to larval development. Timing of cell divisions was inferred from the classical post-embryonic lineage^11^. See Figure S4 for details. **(C)** Exploratory activity, inferred from patterns of locomotion, during the L2, L3, and L4 larval stages. Original data reported by Stern et al.^48^. Dashed grey lines represent displacement over the previous 10 seconds by individual worms over time. Black lines represent population averages. **(D)** Pharyngeal pumping rates in the first and second halves of L2, L3, and L4. Each dot is one individual. *p*-values were calculated using Kolmogorov-Smirnov tests. See Table S1 for sample sizes.

The episodes of environmental sensitivity appear to take place during the latter portions of developmental stages (Figure 4A) and typically involve perception of food. Thus, they may be coupled to the programs that regulate starvation arrests early in all larval stages^38^ and act as checkpoints to prevent development in the absence of food^39^. Two additional points need to be made. First, starvation checkpoints represent regulated decisions made based on sensing the environment^40^, not the inability to continue development due to insufficient resources^39,41^. Second, although in some cases the stage during which environmental sensing occurs is immediately followed by the stage during which development is altered, this need not be the case. For example, food deprivation during early, but not late larval development, can suppress the emergence of male-specific patterns of synaptic connectivity^42^. Critically, although food is a major determinant of developmental trajectories, other environmental signals, including social cues as we showed here, are also important.

The plausible reason for the periods of environmental sensitivity that we envision (Figure 4A) is to gather information relevant to making decisions regarding major developmental events. One such event is molting at the end of each larval stage that involves the shedding of a cuticle^43^. Bursts of cell division events also tend to occur in the beginning of larval stages, particularly L2, L3, and L4 (Figure 4B, S4), and are among the developmental events that are arrested at specific checkpoints if adequate food is absent^39^. Recent studies have demonstrated that expression of several thousand genes oscillates during larval development^44,45^ and that nutrient sensing regulates rhythmic gene expression^46^. We envision that during environmentally sensitive periods in each larval stage worms collect information, certainly on nutrient availability, but as we argue here also on social milieu and probably other features of the environment. These sensory inputs can subsequently be integrated to adjust oscillating gene expression and/or alter the implementation of developmental decisions.

A possible hallmark of environmentally sensitive periods is that they may be enriched for behavioral states^47^ that are less frequent during other periods within a given larval stage. Since nutrient sensing is likely a major feature of environmentally sensitive periods, they may be enriched for food-associated behaviors. A study of *C. elegans* locomotion during larval development^48^ offers support for this hypothesis. On food *C. elegans* larvae engage in interspersed episodes of roaming (high-velocity directional movements with few turns) and dwelling (slower movement with more frequent turns and reversals)^49-52^. Stern *et al*. have found that the early parts of L3 and L4 were dominated by roaming, whereas the latter parts were spent mostly dwelling^48^. Both behavior and developmental processes may be regulated differently during the first larval stage (L1), in which worms show different behavioral profiles under fed conditions^48^ and following starvation^53^. The patterns of oscillating gene expression^45^, growth^6,10^ and development (Figure 4B) in L1 also appear to be different than during other stages. The periods of environmental sensitivity we propose (Figure 4A) correspond well with the periods of reduced locomotion (Figure 4C shows plots of locomotion based on the data from^48^). Patterns of stereotyped movement originate before the onset of larval development, during embryogenesis^54^. In what is either a coincidence or a yet-to-be understood connection, oscillations in gene expression^45^ and stereotyped movement during embryogenesis^54^ are initiated around the same time (∼400 minutes post fertilization), approximately halfway through embryogenesis^13^.

Roaming states are associated with increased locomotion and global exploration, whereas dwelling states with less directional movement and local exploitation of resources^50-52^. Consistent with the idea of coordinated motor programs^55^, we found that during earlier portions of larval stages worms showed significantly lower rates of pharyngeal pumping, a proxy for food intake^56^ (Figure 4D). We interpret the distinct patterns of locomotion and feeding during the periods of environmental sensitivity as an extension of the maximally informative foraging model^57^. At the beginning of larval stages worms pursue global exploration of their environment seeking optimal conditions with respect to food, temperature, osmolarity, social signals, etc. The ticking developmental clock inexorably shifts the strategy toward dwelling, i.e., intense exploitation of local resources. Once identified, the dwelling location will essentially be the site of major developmental events at the beginning of the next larval stage. Therefore, worms convert sensory inputs from the environment into decisions regarding which of the possible developmental trajectories is best suited for the locale. These can range from developmental arrest if the conditions are poor to accelerated development in the presence of potential mates.

It is notable that different behavioral states during early vs. late larval stages could be detected even under the relatively optimized and certainly homogenous laboratory conditions. It may be expected that in patchy, ephemeral environments of the natural *C. elegans* habitats^58^, the distinctions between behavioral patterns in the early vs. late portions of a larval stage would become more pronounced.

### Parallels in other species

Phenomena similar to the ones described above have been documented in other species.

Developmental plasticity is a common feature in multiple animal lineages^59-61^. Consequently, windows of heightened sensitivity to specific environmental signals that direct developmental trajectories and appropriate behaviors must be common as well. It is well known that larvae of different ages, for example in *D. melanogaster*, exhibit different patterns of locomotory behavior^62^. High-resolution long-term observation of animal behavior promises to reveal previously unknown behavioral phases^63^. For example, in Drosophila continuous monitoring of larval crawling behavior on the timescales comparable to the duration of one larval instar revealed apparent epochs of faster directional movement with fewer turns followed by prolonged periods of slower movement with more turns^64^. This pattern is strikingly similar to the one observed during long-term behavioral monitoring of *C. elegans* larvae^48^ (Figure 4A, C).

Drosophila larvae display characteristic patterns of activity associated with cessation of feeding at the end of larval development^65^. Such association between feeding behavior and developmental progression likely reflects the requirement to attain “critical weight” to initiate metamorphosis^66^. Food quantity and quality greatly affects larval development^67,68^. Consequently, developmental stage can shape food choices and behavior^69^. The relationship between growth and development on the one hand and behavioral states on the other may be quite granular as evidenced by the acute inhibition of insulin-producing cells during bouts of locomotor activity in Drosophila^70^. Finally, although centrally important for regulating larval development and behavior, food is just one salient environmental variable (e.g.^71^), with specific roles of others remaining to be discovered.

## Conclusions

Larval stages in *C. elegans* are partitioned into distinct epochs of exploration and consumption as well as discrete windows of heightened sensitivity to the environment, including nutrients and social signals. In this way, behavior matches the execution of developmental programs to the variable and largely unpredictable conditions of the habitat. We expect that in larvae of other species major developmental transitions should be preceded by episodes of intense exploration and sensing to ensure that the environment is suitable for these costly commitments. It would be interesting to identify which environmental signals, other than food availability, are particularly relevant for the progression of the immediately following developmental events in different larval stages. It is likely that, just like in *C. elegans*, in all species instead of uniform behavior in each larval stage, there exist distinct epochs enriched for different internal states that extend beyond the locomotion and eating behaviors addressed here. Testing these predictions promises to enrich understanding of how behaviors are structured along the life history continuum and reveal the mechanisms that coordinate behavior and development.

## Acknowledgements

We are grateful to Rick Morimoto for generous hospitality. We thank David Sherwood for NK881, Judith Kimble for the Anti-SP56 antibody, Shay Stern for sharing data and comments, and Marco Gallio and Dan Tracey for comments and advice. This work was funded in part by NSF (IOS-1755244) and NIH (R01GM126125) grants to IR. We thank WormBase and the Caenorhabditis Genetics Center (CGC).

WormBase is supported by grant U41 HG002223 from the National Human Genome Research Institute at the NIH, the UK Medical Research Council, and the UK Biotechnology and Biological Sciences Research Council. The CGC is funded by the NIH Office of Research Infrastructure Programs (P40 OD010440).

## Materials and Methods

### *C. elegans* handling and strains

*C. elegans* nematodes were maintained using standard methods^72^ on NGM plates seeded with OP50 *E. coli*. Unless otherwise noted, all experiments were done at 20°C with synchronized N2 hermaphrodites grown in isolation (i.e., 1 worm/plate) and experiments were conducted with paired controls. Synchronous cultures were obtained by treating populations of gravid adults with an alkaline hypochlorite solution and hatching embryos overnight (≤ 16 hours) in M9 buffer with rotation^73^. Arrested L1 larvae were plated onto 60-mm lawn plates of OP50 at a density of 20-30 worms/plate and the time of plating was designated as the release from L1 arrest. All worm ages in this study are expressed as hours following release from the L1 arrest. Immediately after, worms were singled onto 35-mm plates that were seeded with 5μl of 1:10 OP50 dilution and grown overnight. Male-conditioned plates (MCPs) were made by placing a single young adult N2 male for 24 hours and removing it prior to the start of the experiments. The following strains were used: N2 wild type, SV1009 heIs63 [*wrt-2*p::GFP::PH + *wrt-2*p::GFP::H2B + *lin-48*p::mCherry]^74^, PD4667 ayIs7 [*hlh-8*::GFP fusion + *dpy-20*(+)]^75^, NK881 [qyIs166[*cdh-3*>GFP::CAAX]; qyIs127[*lam-1*::mCherry]^76^.

### Imaging progression of developmental events

For all imaging, with the exception of body volume, worms were mounted onto 2% agarose slides, observed, and imaged on a Leica DM5000B microscope using a Retiga 2000R camera.

L4 vulva substages (Figure 1B) were scored manually based on the morphology of the vulval lumen at 50 hours. Standard definitions^77,78^ were adapted to the specifics of our experimental procedures to derive a consistent series of substages. L4.0 could be distinguished from the late L3 by observing the shed cuticle. L4.1 = Vulva undergoes invagination. L4.2 = Invagination progresses beyond the ventral P6.p great-granddaughter cells. L4.3 = Convex sides develop. L4.4 = Upside-down T shape with sharp corners forms. L4.5 = Rounded corners and “fingers” develop. L4.6 = “Fingers” start pointing ventrally. L4.7 = Vulva starts collapsing forming a maple leaf shape. L4.8 = Vulval collapse progresses leaving a small invaginated space. L4.9 = Vulval lips protrude outwards but remain covered by the cuticle. See Figure S1A for representative images.

To quantify the extent of germline development (Figure 1C, D), worms were scored for the number of oocytes completely spanning the gonad lumen and for the number of fertilized embryos in the uterus.

To estimate body volume (Figure 1F, G) worms were imaged without removal from 35-mm NGM plates. The images were taken on an Olympus SZ61 stereomicroscope fitted with a Lumenera Infinity 2 camera and processed manually using ImageJ. Length was measured using the segmented line tool by skeletonizing the worm following the midline from most anterior to most posterior discernable points. Width was measured at the level of the vulva, if identifiable, or at ∼2/3 of body length. The image in Figure 1E shows skeletonization used for ImageJ processing.

Progression of certain developmental events in the soma was ascertained using reporter strains. Seam cell development (Figure 2A) was monitored using the SV1009 strain (*wrt-2*::GFP) that helped to visualize divided and adhered cells. See Figure S1B for representative images. Sex myoblast divisions (Figure 2B) were scored in the PD4667 strain (*hlh-8*::GFP). See Figure S1C for representative images used for staging. Anchor cell (AC) invasion (Figure 2C) was scored in the NK881 strain (*cdh-3*::GFP).

Staging (see Figure S1D for representative images) was based primarily on the shape of the cell’s ventral side. AC1 = long curving ventral side not attached to P6 daughter cells. AC2 = flat and short ventral side indicates attachment. AC3 = invasive protrusion forms a V-shape. AC4 = invasive protrusion retracts forming an M-shape.

Gonad length was measured by following the midline of the gonad using the segmented line tool in ImageJ if the entirety of the gonad was visible. Turning phases were defined as follows: TP1 = gonad and distal tip cell (DTC) are extending along the ventral side away from the vulva; TP1/2 = gonad is continuing to extend, DTC has initiated a dorsal turn; TP2 = gonad and DTC extend dorsally; TP2/3 = DTC begins to extend along the dorsal side toward the midline defined by the vulva. See Figure S1E for representative images.

For counting nuclei in the Progenitor Zone (for definition, see^79^), hermaphrodites were stained with DAPI (4′,6-diamidino-2-phenylindole) using a previously described^2^ variation of a published protocol^80^. In addition to mitotic nuclei, this population contains some nuclei in the early stages of meiosis^81^. For sperm counts, 54-hour hermaphrodites were stained with DAPI as above.

### Immunohistochemistry

Worm dissection and antibody staining were modified from a published protocol^82^. At 42, 44, and 46 hours control and MCP hermaphrodites were picked into 30μL PBS-0.1% Tween 20 with 0.25mM levamisole in the bottom half of a large glass petri dish. Hermaphrodites were cut with a scalpel to extrude the germline. ∼30 animals per condition were dissected in ∼5 minutes. The dissected animals were transferred to a 1.5mL microcentrifuge tube and incubated with 3% paraformaldehyde in PBS-Tween for 30 minutes at 20°C with rocking. The paraformaldehyde was washed off and the worms were fixed in -20°C methanol overnight. The methanol was washed off and the worms were blocked with 3% bovine serum albumin in PBS-Tween for 30 minutes at 20°C with rocking. The blocking agent was washed off and the worms were incubated overnight with the primary antibody (Anti-SP56 antibody^21^, a gift from the Kimble lab, diluted 1:50 in block) at 4°C with rocking. The following morning the worms were washed 3X with PBS-Tween at 20°C with rocking (≥ 10 minutes) and subsequently incubated with the secondary antibody (Goat Anti-Mouse IgG H&L Alexa Fluor® 488 Abcam150113, diluted 1:1000 in PBS) for two hours at 20°C with rocking. The worms were washed again 3X with PBS-Tween as above, suspended in 12μL Vectashield with DAPI, and transferred to 2% agarose pads. Imaging was performed as above.

### Scoring pumping at the L3-to-L4 transition

At 32 hours singled larvae were removed from the 20°C incubator and continuously observed until all observed worms entered L4 (39 hours, at room temperature). For each worm, grinder movement was observed for 10 seconds every 20 minutes on a Leica MZ16 stereomicroscope. Worms were categorized as L3 if active pumping was observed, as quiescent if no pumping was observed, or L4 if pumping and an invaginated vulva were observed. To limit bias, we scored five worms from control plates, followed by five MCP worms, until all worms were examined.

### Obtaining the density plot of timing of post-embryonic cell divisions

The unlabeled *C. elegans* lineage diagram was obtained from WormAtlas (https://www.wormatlas.org/images/lineage.png) and cropped to include only post-embryonic divisions (final size: 1034 × 2317 height and width, respectively). The image consisted of 3 types of elements: vertical lines with width = 2 pixels (px) and height >> 2px, horizontal lines with width of >> 2px and height = 2px, and “X” markers consisting of 5 elements with height and width of each ≤ 2px. Since horizontal lines represented division events, we used a Python script to scan the image top-down by row (increasing Y coordinate, aligned with time progression) and deleted all elements with width ≤ 2px. The processed image was then scanned by row again to count the number of contiguous regions of black pixels (i.e., discrete horizontal bars), each representing a division event. Sharp peaks appearing at coordinates 448-449 and 684-685 corresponded to Seam Cell divisions. These events occur shortly after the onset of L2 and L3 at ∼16 and ∼25 hours post hatching, respectively. The knowledge of this timing allowed us to convert the time scale from the image Y coordinate (row) to hours of development. Division density was then calculated using an inverse kernel: for value *V*_*i*_ density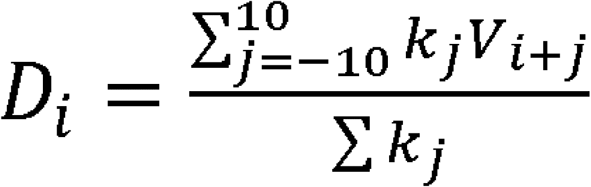 where weight 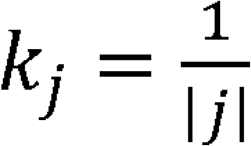, unless *j* =0 when *k* =1. The procedure is shown schematically in Figure S4. The code used to process the post-embryonic lineage diagram was deposited: https://github.com/denisfaer/Faerberg_et_al_2023_Acceleration/blob/main/plotscan.py

### Generating activity profiles

The original data on locomotion of 125 N2 worms throughout larval development were collected by Stern et al.^48^. Durations of larval stages inferred from these data were reported^5^. For each individual’s larval stage *j* activity profile point 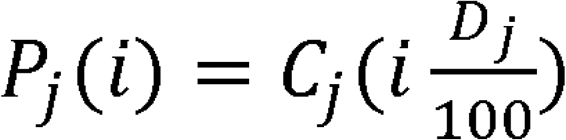 where *i* is the percentage of stage completion, *D*_*j*_ is the duration of the individual’s stage *j*, and *C*_*j*_ (*t*) is the individual’s activity curve as inferred^5^ and cropped to larval stage *j*. The average of P (i) for all individuals for each *i* yielded the population averages plotted in Figure 4C. The code used to generate these profiles was deposited: https://github.com/denisfaer/Faerberg_et_al_2023_Acceleration/blob/main/stageact.pas

### Quantifying the pharyngeal pumping rate

At 19 (first half of L2), 23 (second half of L2), 28 (first half of L3), 34 (second half of L3), 41 (first half of L4) and 46 (second half of L4) hours 50 worms reared on control plates were recorded for 10 seconds on a Leica MZ16 stereomicroscope. For each worm, the number of grinder movements was manually counted and normalized by the duration of the recording in seconds. Worms with no observed contractions or those located off the bacterial lawn were excluded from the analysis.

### Statistical analyses

Significance between population fractions (e.g. Figures 1A, 2A, 3E) was computed in R using the test of equal or given proportions. Significance between quantitative traits (e.g. Figures 1F, 1G, 2F, 4D) was computed in R using the Kolmogorov-Smirnov test. Significance between categorical distributions (e.g. Figures 2B, 2C, 2E) was computed using the chi-square test. Results of all tests are presented in Table S1.

