## Supplemental Figures for "Periods of environmental sensitivity couple larval behavior and development"

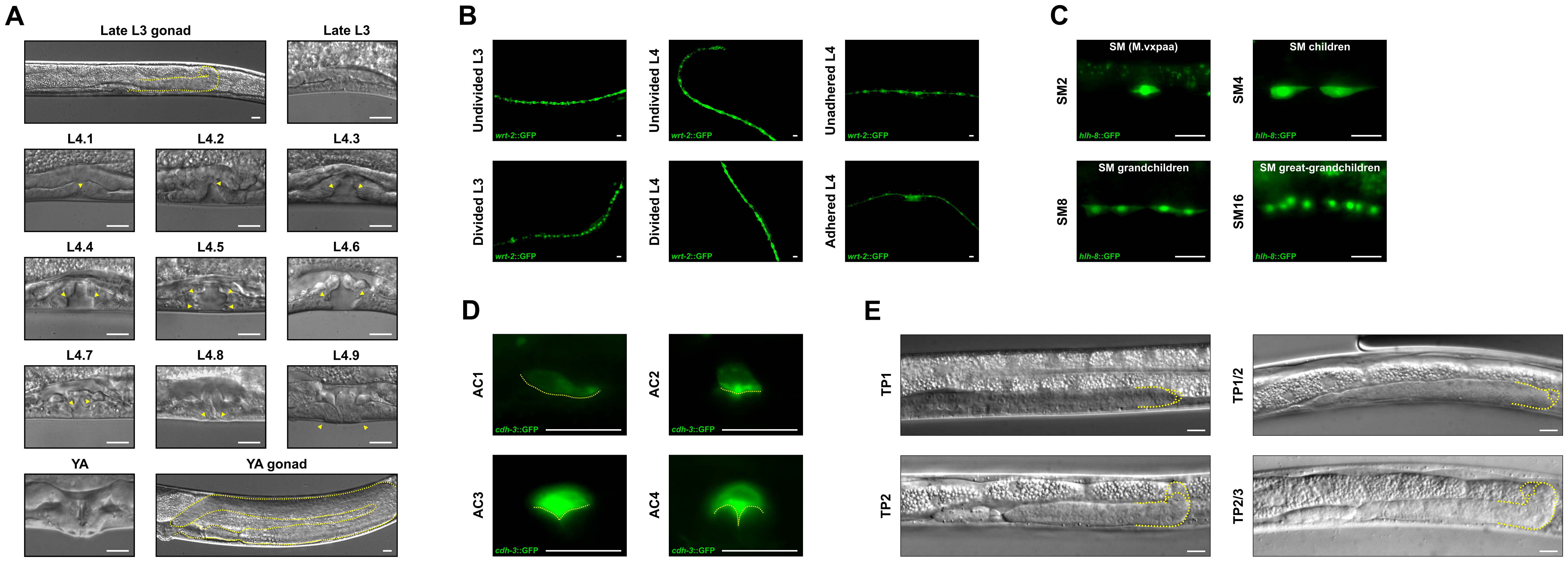
A
Late L3 gonad
Late L3
L4.1
L4.2
L4.3
L4.4
L4.5
L4.6
L4.7
L4.8
L4.9
YA
YA gonad
B
Undivided L3
Undivided L4
Unadhered L4
Divided L3
Divided L4
Adhered L4
wrt-2::GFP
C
SM (M.vxpaa)
SM children
SM grandchildren
SM great-grandchildren
SM2
SM4
SM8
SM16
hlh-8::GFP
D
AC1
AC2
AC3
AC4
cdh-3::GFP
E
TP1
TP1/2
TP2
TP2/3

**Figure S1. Morphological definition of scoring criteria. Related to Figures 1 and 2.**

**(A)** Morphological features of the vulva and the gonad during different substages in late hermaphrodite larvae. Based on previously described definitions[77,78] and adapted to our experimental procedures. See Materials and Methods for stage descriptions. Dashed yellow outlines show gonad progression, yellow arrowheads point to key morphological features. **(B)** Fluorescence microscopy images of seam cells (*wrt-2*::GFP) in representative individuals before (top) and after (bottom) undergoing early-L3 division (left), early-L4 division (middle) or mid-L4 adhesion (right). **(C)** Fluorescence microscopy images of sex myoblasts (*hlh-8*::GFP) undergoing three rounds of divisions. **(D)** Fluorescence microscopy images of the invading anchor cell (*cdh-3*::GFP). See Materials and Methods for stage descriptions. **(E)** Representative DIC images of mid- to late-L3 individuals that show the phases of gonad turning. See Materials and Methods for stage descriptions. Dashed yellow outlines show the distal portion of the gonad and DTC orientation. All scale bars = 10μm.

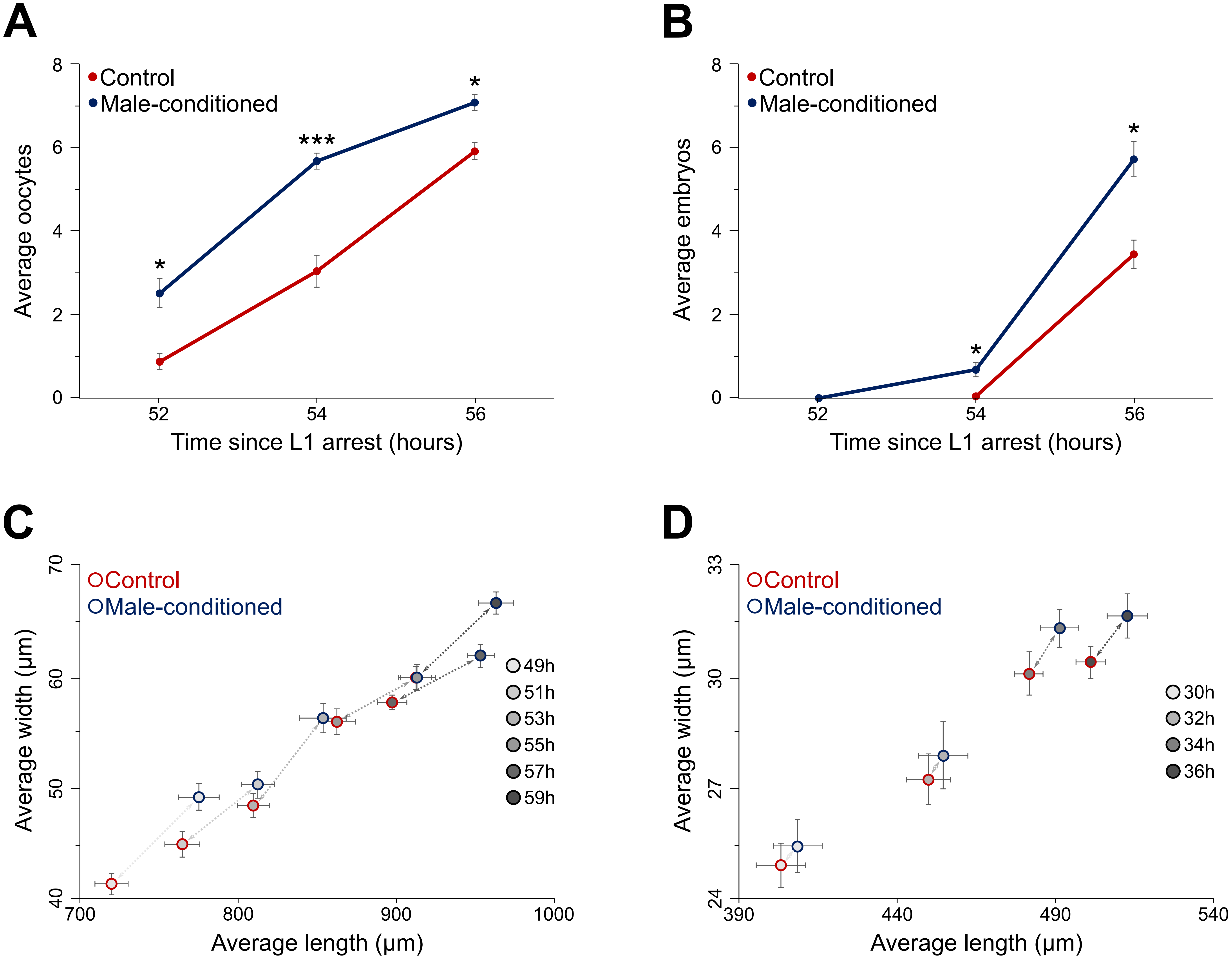
A
Control
Male-conditioned
Average oocytes
Time since L1 arrest (hours)
B
Control
Male-conditioned
Average embryos
Time since L1 arrest (hours)
C
Control
Male-conditioned
49h
51h
53h
55h
57h
59h
Average width (µm)
Average length (µm)
D
Control
Male-conditioned
30h
32h
34h
36h
Average width (µm)
Average length (µm)

**Figure S2. Male-excreted signals accelerate development and growth of hermaphrodite larvae. Related to Figure 1.**

**(A)** Related to Figure 1C. Average number of oocytes spanning the gonad lumen. Whiskers show standard error. **(B)** Related to Figure 1D. Average number of embryos in the uterus. **(C)** Related to Figure 1F. Average length and width of worms emerging into adulthood. Circle color indicates growth condition, circle fill color indicates age, dashed arrows connect the same timepoint in control and MCP groups. Whiskers show standard error. **(D)** Related to Figure 1G. Average length and width of worms in the second half of the L3 stage. Symbol meaning is as in Figure S2C. See Table S1 for sample sizes and statistical analyses.

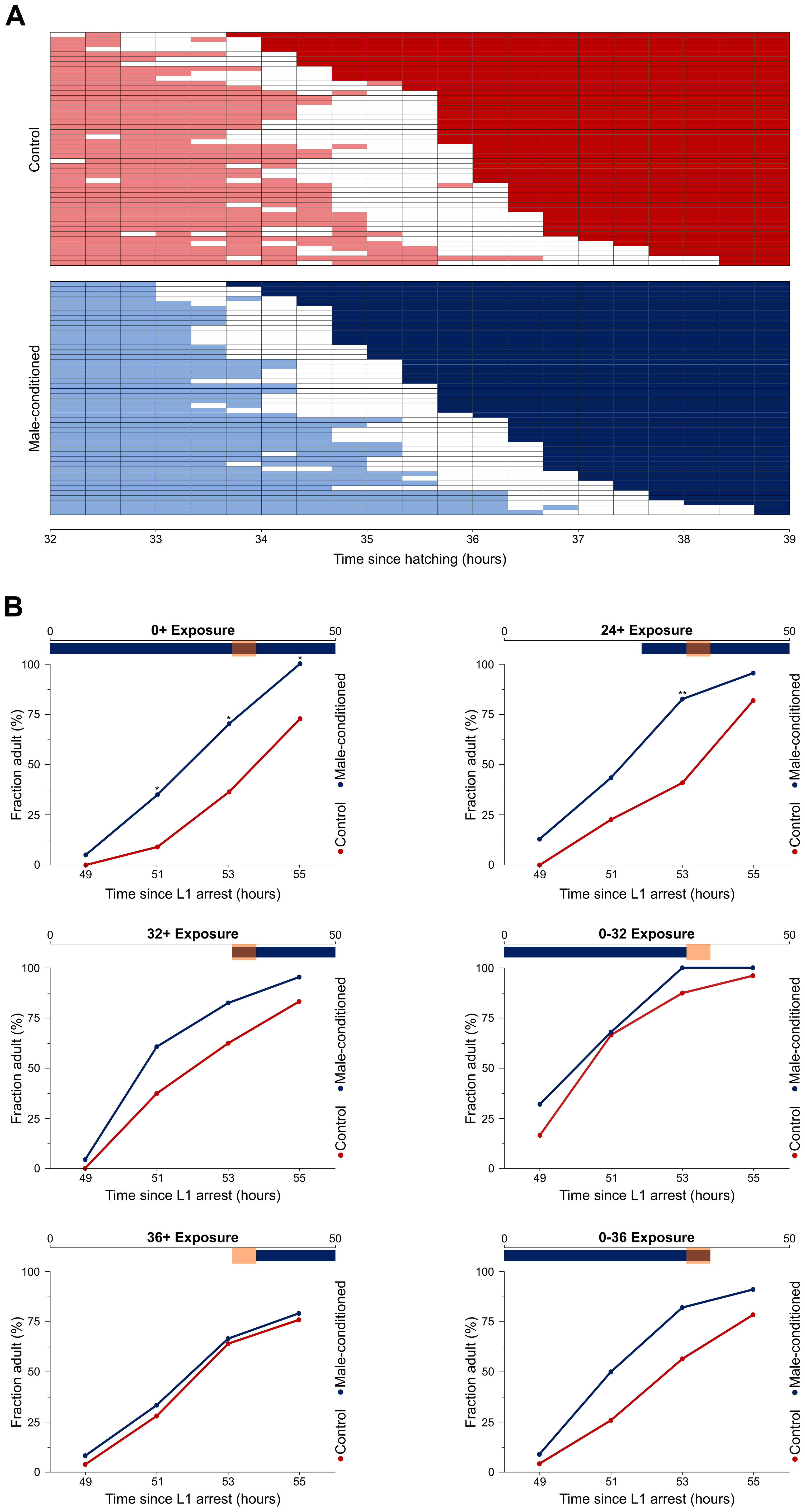

A
Control
Male-conditioned
32
33
34
35
36
37
38
39
Time since hatching (hours)
B
0+ Exposure
24+ Exposure
32+ Exposure
0-32 Exposure
36+ Exposure
0-36 Exposure
0
50
100
75
50
25
0
Fraction adult (%)
49
51
53
55
Time since L1 arrest (hours)
Male-conditioned
Control
*
**

**Figure S3. Male-excreted signals and developmental timing. Related to Figure 3.**

**(A)** The data presented in Figure 3A rearranged in order of the onset of L4. This view highlights the "uncertainty" of lethargus commitment in control worms. **(B)** Representative results of experiments summarized in Figure 3F. Period of exposure to the male-excreted signals is indicated on top of each sub-panel with a blue and numerical notation. Orange box corresponds to the period between 32 and 36 hours. See Table S1 for sample sizes.

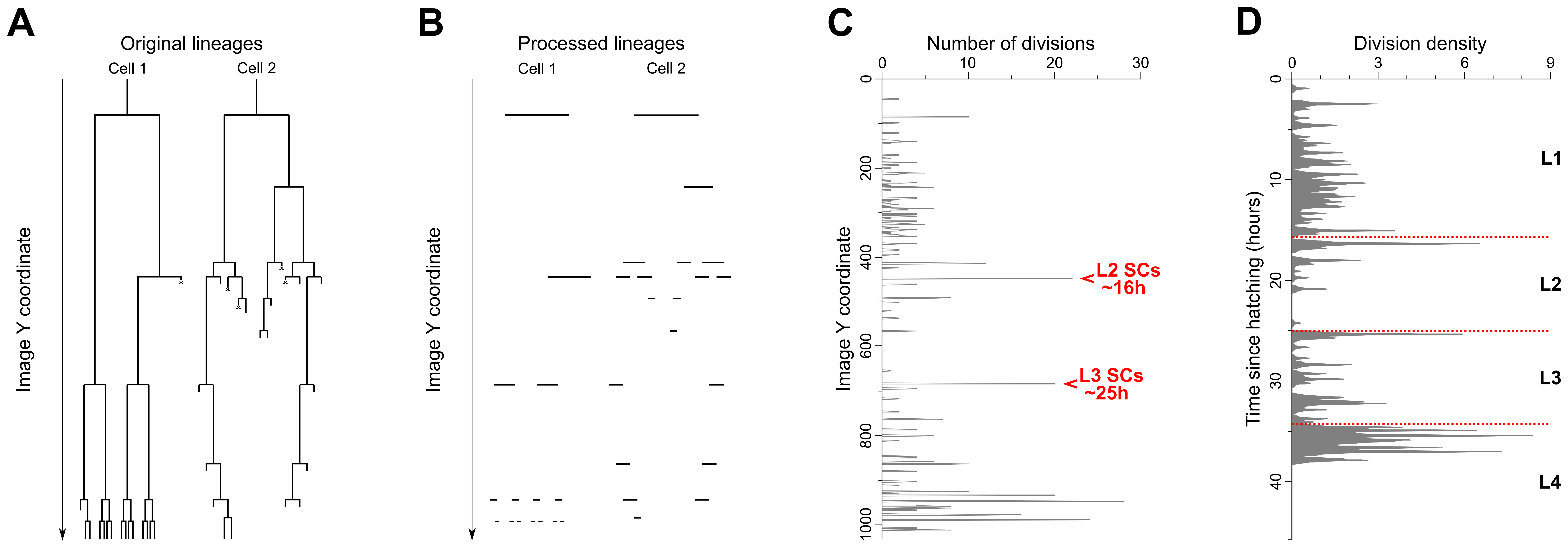

A
Original lineages
Cell 1
Cell 2
Image Y coordinate
B
Processed lineages
Cell 1
Cell 2
Image Y coordinate
C
Number of divisions
0
10
20
30
Image Y coordinate
0
200
400
600
800
1000
L2 SCs
~16h
L3 SCs
~25h
D
Division density
0
3
6
9
Time since hatching (hours)
0
10
20
30
40
L1
L2
L3
L4

**Figure S4. Obtaining the density plot of timing of post-embryonic cell divisions. Related to Figure 4.**

**(A)** A hypothetical lineage tree similar to the post-embryonic lineage diagram of *C. elegans*. **(B)** Image processing to infer division times in the tree shown in panel A. Processing removed all objects in every row with width ≤ 2 pixels, but retained horizontal lines representing divisions. Removed items included vertical lines and "X" markers representing cell deaths. Scanning each row and counting contiguous blocks of black pixels yields the number of cell division events. Panels A and B are shown for the purpose of illustrating the approach. **(C)** Unlike the hypothetical schematics in panels A and B, this panel shows the analysis of actual division events during *C. elegans* post-embryonic development as given in the classical lineage diagram[11] (https://www.wormatlas.org/images/lineage.png). Inferred number of divisions (calculated as described in B) at each timepoint (the Y axis denotes time). Marked spikes in rows ~450 and ~680 are due to L2 and L3 Seam Cell divisions occurring at ~16 and ~25 hours post hatching. These landmarks allow a conversion from a time scale expressed in "rows" to a time scale in absolute hours of development. **(D)** A kernel density plot of the data from panel C visually represents some inter-individual variability in division timing. The time scale was converted into hours, as described above. Dashed red lines demarcate boundaries between larval stages. More details in Materials and Methods.
